# Stiffness regulates dendritic cell and macrophage subtype development and increased stiffness induces a tumor-associated macrophage phenotype in cancer co-cultures

**DOI:** 10.1101/2024.03.11.584525

**Authors:** Carla Guenther

## Abstract

Mechanical properties of tissues including their stiffness change throughout our lives, during both healthy development but also during chronic diseases like cancer (1-4). How changes to stiffness, occurring during cancer progression, impact leukocytes is unknown. To address this, myeloid phenotypes resulting from mono- and cancer co-cultures of primary murine and human myeloid cells on 2D and 3D hydrogels with varying stiffnesses were analyzed. On soft hydrogels, conventional DCs (cDCs) developed, whereas on stiff hydrogels plasmacytoid DCs (pDCs) developed. Cell populations expressing macrophage markers CD14, Ly6C, and CD16 also increased on stiff hydrogels. In cancer co-cultures, CD86^+^ populations decreased on higher stiffnesses across four different cancer types. High stiffness also led to increased vascular endothelial growth factor A (VEGFA), matrix metalloproteinases (MMP) and CD206 expression; ‘M2’ markers expressed by tumour-associated macrophages (TAMs) (5). Indeed, the majority of CD11c^+^ cells expressed CD206 across human cancer models. Targeting the PI3K/Akt pathway led to a decrease in CD206^+^ cells in murine cultures only, while human CD86^+^ cells increased.

Increased stiffness in cancer could, thus, lead to the dysregulation of infiltrating myeloid cells and shift their phenotypes towards a M2-like TAM phenotype, thereby actively enabling tumor progression. Additionally, stiffness-dependent signaling appears species-dependent, potentially contributing to the high failure rate of clinical trials (6).

## Introduction

Most of our knowledge on the immune system is based on chemical stimulation, such as from pathogens (pathogen associated molecular patterns, PAMPs), signaling molecules or parts of damaged cells (damage associated molecular patterns, DAMPs). However, all cell activity happens within microenvironments which present different mechanical information, such as stiffness (the extent to which materials resist deformation) and stiffness changes as a result of chronical diseases such as cancer (4). In addition, chronic cell damage and the release of DAMPs may contribute to cancer development (7), while DAMP levels increase in advanced tumors resulting from chemotherapy (8). Both DAMPs and PAMPs are recognized by C-type lectin receptors (CLRs) such as Mincle and Dectin-1 (9). However, how mechanical information affects our immune system, specifically in terms of CLR signaling, remains poorly understood.

Dendritic cells (DCs) and macrophages are central components of the immune system. These innate immune cells survey various tissues and sense abnormal ligands such as PAMPs and DAMPs via receptors such as CLRs. DCs then take up sensed antigens via adhesion receptors such as integrins during phagocytosis and migrate to the nearest lymph node to present the antigen to T-cells, which initiates the adaptive immune response. Not only are integrins used for phagocytosis, but also for sensing and relaying mechanical information into the cell in order to initiate cell responses (10). One example of this is how cells typically migrate towards stiffer substrates; a process known as durotaxis (11).

The aim of this study was to elucidate how differences in tissue stiffness impact DC development. Here, it was shown that stiffness regulates DC and macrophage phenotypes beyond maturity, polarizing the mature phenotype into cDCs or pDCs. While cDCs developed more on softer substrates, high stiffnesses induced pDC development and increased CD14^+^ cell populations along with the expression of other macrophage markers. Moreover, 3D monocultures in hydrogels with different stiffnesses showed a similar tendency to develop these phenotypes. These phenotypes developed due to stiffness signals alone and also occurred and became more pronounced following cell stimulation with PAMP and DAMP CLR ligands. These phenotypes were mediated by transcription factors such as Ikaros as well as STAT5 and STAT3 signaling. Stiffness-dependent cDC and pDC polarization also emerged in murine 3D co-cultures with melanoma cells, while both populations decreased in colon and breast cancer co-cultures. Instead, high stiffnesses induced CD14^+^ populations across all three cancer types. In human cancer co-cultures, most CD11c^+^ cells expressed CD206 on higher stiffnesses and high stiffness induced expression and production of VEGFA and MMPs in mice and humans. This suggests that stiffness contributes to switching myeloid cells towards a TAM phenotype.

Treating cell cultures with Akt Inhibitor IV reduced CD206 expression in murine CD11c^+^ cells, but not in human cells. Instead, inhibiting Akt signaling in human CD11c^+^ cells led to an increase of CD86 expression, but not in murine cells. This indicates that Akt is a central regulator of the myeloid cell phenotype, but its specific function is species-dependent. Nevertheless, these data suggest targeting Akt in tumor-associated myeloid cells might be a valid strategy to switch an anti-inflammatory to a pro-inflammatory tumor microenvironment to induce antitumor responses and improve patient survival.

## Results

### Substrate stiffness polarizes DC subtype phenotype into cDC vs pDC in 2D and 3D

To investigate the stiffness impact on DC phenotype, bone marrow (BM) cells were differentiated into DCs using granulocyte macrophage colony stimulating factor (GMCSF). Cells were cultured on commercially available silicone hydrogels with different stiffnesses (0.2 kPa, 0.5 kPa, 2 kPa, 8 kPa, 16 kPa, 32 kPa and 64 kPa which roughly correlate with different tissues (e.g. brain, lung, kidney, muscle, eye and bone). As a positive control conventional tissue culture plastic was used. To present the same chemical surface to the cells, hydrogels and plastic were coated with 100 μg/ml collagen. This culture system resulted in at least 60% CD11c^+^ cell populations across all stiffnesses (Supplementary Figure 1a). Importantly, different DC phenotypes were observed across the different stiffnesses even without further stimulation, with clear populations emerging as shown in Figure 1a when analyzed via flow cytometry (gating strategy shown in Supplementary Figure 1b). On stiff substrates percentages of CD317^+^, Siglec-H^+^, CD319^+^ and CD205^+^ cells increased significantly (Figure 1b), while B220 expression level showed a tendency to be increased (Supplementary Figure 1c) and TNF production increased, all of which associated with the pDC phenotype (Figure 1b). There was a significant increase in TNF production, from 0.2 to 0.5 kPa, indicating that even extremely small stiffness changes can significantly impact the DC phenotype. Additionally, percentages of CD14^+^ and CD16^+^ cells were increased (Figure 1c) on stiffer substrates. On soft substrates, CD80^+^, CD86^+^ and Ikaros^+^ cell populations and IL-6 production were increased (Figure 1d), which is associated with cDCs and decreased integrin signaling (12, 13).

**Figure 1:**
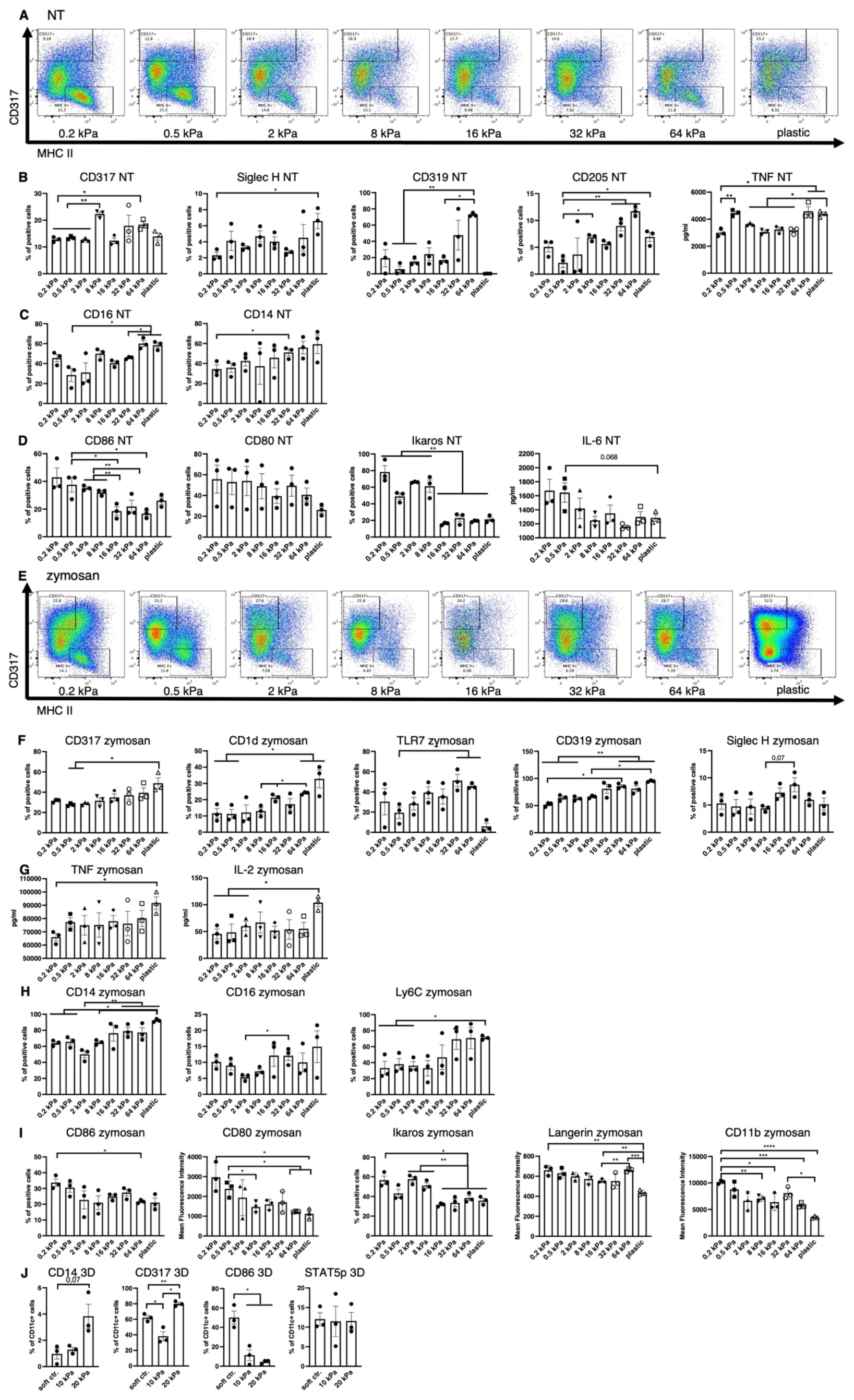
Low stiffness induces the cDC and high stiffness the pDC phenotype. **A)** An example of flow plots depicting the shifting CD317^+^ and MHCII^+^ cell populations across the 8 stiffnesses that arose in BM derived cell cultures without stimulation (NT, not treated). **B)** Cell populations positive for pDC associated surface markers were measured in resting cells (NT, not treated) via flow cytometry. Percentages of all living cells are shown. The production of TNF was measured via ELISA. **C)** Cell populations positive for macrophage markers were measured in resting cells (NT, not treated) via flow cytometry. Percentages of all living cells are shown. **D)** Soft substrates induced an increase of cell populations positive for cDC associated markers as well as IL-6 production in resting cells. Cell populations positive for cDC-associated surface markers were measured in resting cells (NT, not treated) via flow cytometry. Percentages of all living cells are shown. The production of IL-6 was measured via ELISA. **E)** An example of flow plots depicting the shifting CD317^+^ and MHCII^+^ cell populations across the 8 stiffnesses that arose in BM-derived cell cultures after zymosan stimulation (24 h). **F)** Cell populations positive for pDC associated surface markers were measured in zymosan-stimulated (24 h) cells via flow cytometry. Percentages of all living cells are shown. **G)** The production of TNF and IL-2 was measured via ELISA. **H)** Cell populations positive for macrophage surface markers were measured in zymosan-stimulated (24 h) cells via flow cytometry. Percentages of all living cells are shown. **I)** Cell populations positive for cDC associated surface markers were measured in zymosan-stimulated (24 h) cells via flow cytometry. Percentages of all living cells are shown for Langerin and CD11b expression levels (MFIs) are shown. **J)** The cell populations of BM-derived cells cultured in 3D alginate/collagen hydrogels (25). Surface marker expression was measured via flow cytometry. Cell populations were first gated for CD11c in order to exclude gel particles and the graphs depict the percentage of CD11c^+^ cells. 10 kPa and 20 kPa were stiffened by addition of CaCl_2_ and soft controls were not stiffened (soft ctrl.). **A–J)** All experiments were performed with three biological repeats per technical repeat and at least two technical repeats in total. One representative technical repeat is shown in each graph and statistics are based on three biological repeats per condition depicted in the graph. All p values are shown: *p < 0.05; **p < 0.01; ***p < 0.005; ****p < 0.0001. All error bars are shown as SEM.

Following stimulation with zymosan, these phenotypes did not disappear (Figure 1e). A seemingly linear increase in CD317^+^ cells as well as an increase of CD1d^+^, TLR7^+^, CD319^+^ and Siglec-H^+^ (trend, not significant) populations (Figure 1f) was observed on higher stiffnesses, along with an increased production of IL-2 and TNF (Figure 1g). Stiffer substrates also gave rise to increased populations of cells positive for the macrophage markers CD14, CD16, and Ly6C (Figure 1h). On soft substrates CD86^+^ and Ikaros^+^ cell populations increased while CD80, Langerin and CD11b expression levels also increased (Figure 1i). Furthermore, these phenotypes arose regardless of the ligand used for stimulation (DAMP or PAMP), as shown for the example of CD86 and CD80 (Supplementary Figure 1d).

To validate these results in a more *in vivo–*like model, bone marrow derived DC (BMDC) monocultures in 3D alginate/collagen hydrogels were done, resulting in similar decreases in CD11c^+^/CD86^+^ cells on high stiffnesses, while CD11c^+^/CD317^+^ and CD11c^+^/CD14^+^ cell populations increased on high stiffnesses (Figure 1j). There was no difference between CD11c^+^/STAT5p^+^ populations across stiffnesses.

To identify the central regulators governing these phenotypical changes, RNA sequencing was performed on 2D cultures derived from the two most extreme stiffnesses, 0.2 kPa and plastic both with and without zymosan stimulation. Beyond the primary zymosan effect, there was also a smaller set of genes persistently upregulated on soft substrates regardless of zymosan stimulation (Figure 2a). Upon further analysis, multiple interferon-associated genes were discovered to be upregulated on plastic as well as several transcription factors, such as *Etv3, Etv6*, and *Il4ra* (Figure 2b). On 0.2 kPa gels, multiple genes associated with cDCs were upregulated, such as *il12r, il12rb2, Cd1d1, Marco*, and *Ptprn* (Figure 2c). As DC phenotypes are regulated by signal transducer and activator of transcription (STAT) signaling (14), expression of these transcription factors was also analyzed (Figure 2d). While zymosan treatment led to *Stat1* and *Stat2* upregulation compared to unstimulated controls, there was also a significant increase on plastic compared to 0.2 kPa gels for both genes (Figures 2d+e). This pattern was also observed for *Stat3* and *Stat5a*, visible in the heatmap (Figures 2d+e). Notably, *Stat4* was upregulated on 0.2 kPa and downregulated on plastic.

**Figure 2:**
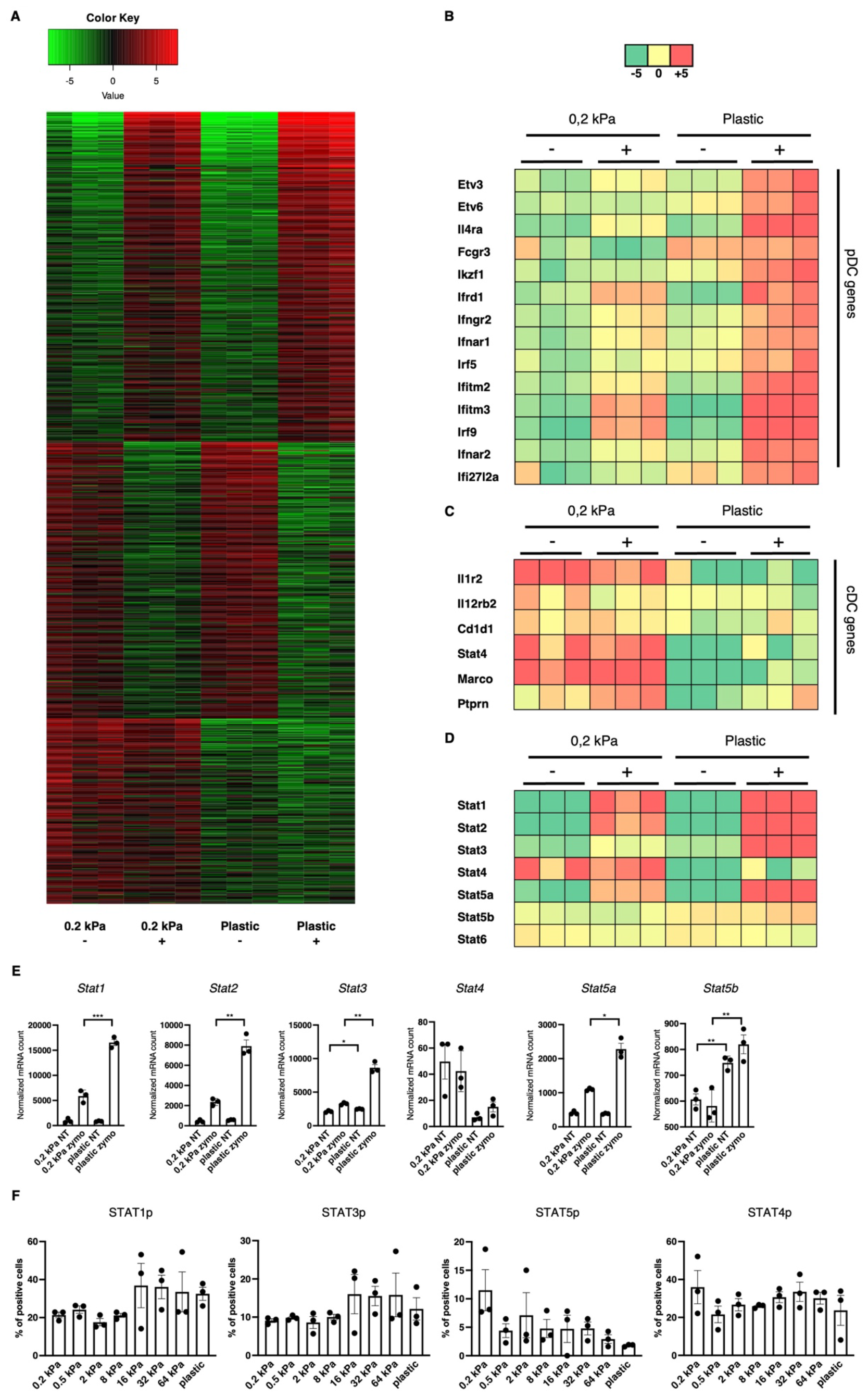
Stiffness-induced cDC and pDC phenotypes are mediated by STAT signaling. **A)** Heatmap depicting global gene expression changes between DCs cultured on 0.2 kPa hydrogels (+/-zymosan) and DCs cultured on plastic (+/-zymosan). Zymosan stimulation was done for 20 h. **B)** Genes associated with the pDC phenotype were upregulated on plastic, **C)** while genes associated with the cDC phenotype were upregulated on 0.2 kPa hydrogels. **D)** Heatmap depicting STAT gene expression changes. **E)** Bar graphs depicting normalized mRNA counts of STAT genes. **F)** Bar graphs depicting STAT phosphorylation at the end of BMDC cultures measured via flow cytometry. The percentages for all living cells are shown. RNAseq experiments were performed using three biological repeats per condition. Flow cytometry experiments were performed with three biological repeats per technical repeat and at least two technical repeats in total. One representative technical repeat is shown in each graph. Three biological repeats per condition were used for the statistical analysis. All p values are shown: *p < 0.05; **p < 0.01. All error bars are shown as SEM.

STAT signaling is predominantly mediated by phosphorylation (15). Thus, to confirm the RNA results, we investigated STAT phosphorylation via flow cytometry (Figure 2f). While no significant differences were observed, several trends emerged. First, STAT1p and STAT3p were predominantly elevated on stiffer substrates, suggesting their involvement in mediating the pDC phenotype, mirroring previous reports (14, 16). However, Stat5 phosphorylation decreased on stiffer substrates. While the cDC phenotype has been shown to be mediated by Ikaros (Figures 1c-f) (13), others studies indicated that STAT5 suppresses the pDC phenotype (17). Lastly, STAT4 phosphorylation did not mirror the RNA expression data, as no consistent stiffness dependent effect on STAT4p could be observed. It is believed there is currently no established role for STAT4 in DC development, however Remoli et al. reported STAT4’s interferon 1– dependent presence in mature DCs (18), which is in line with the results presented here. It is important to note that here, STAT phosphorylation was analyzed at the end of the culture and these results represent a baseline in these cells.

### High stiffness induces a macrophage phenotype in 3D cancer co-cultures

During cancer progression, immune cells including DCs frequently become dysregulated and the stiffness of the tissue changes, especially due to stromal changes in extracellular matrix (ECM) production (19). Melanoma specifically has been shown to increase skin stiffness, from 2–3 kPa to around 25 kPa (4). cDCs as well as pDCs have been found to be beneficial for anti-tumor responses to breast cancer (20), while CD14^+^ cells are associated with immunosuppression across cancer types (21). It was thus hypothesized that a tumor-associated stiffness increase contributes to the dysregulation of innate immune cells such as DCs, macrophages, and their progenitors. In tumor progression, many molecules are produced, some of which are directly responsible for the stiffness of tumors, such as the tumor ECM (19). To test if the tumor ECM overrides the stiffness effect, Matrigel, a commercially available tumor-derived basement membrane, was used to coat hydrogels and these were compared with collagen-coated hydrogels. Steady-state cultures resulted in no significant changes in populations, as shown in the CD86^+^, CD205^+^, CD317^+^ and CD14^+^ cell populations (Figure 3a). This indicates that tumor-derived ECM molecules do not alter the stiffness-mediated DC and macrophage phenotypes.

**Figure 3.**
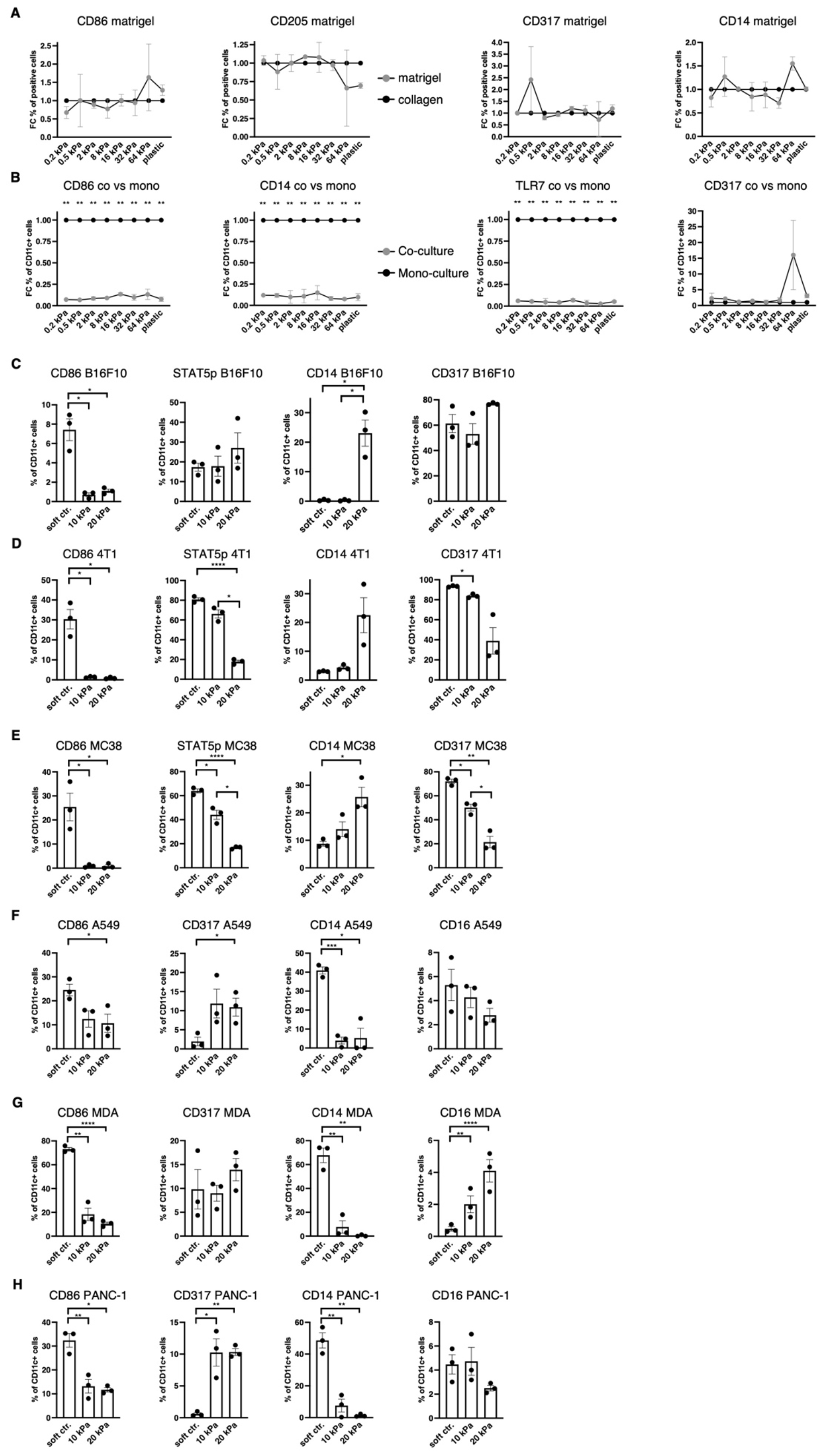
Increased stiffness in cancer co-cultures contributes to cDC and pDC phenotype dysregulation. **A+B)** For all graphs, individual control (collagen or monoculture) percentages were set as 1 for each stiffness, upon which the corresponding fold change was calculated. **A)** Surface marker expression of resting BMDC cultures on collagen or matrigel measured via flow cytometry. The percentages of all living cells are shown. **B)** Surface marker expression of resting BMDC cultures compared to cancer co-cultures (B16F10). Cell percentages are based on CD11c^+^ cell populations and monocultures were set as a baseline. **C)** Cell percentages of CD11c^+^ cell populations from B16F10 co-cultures. **D)** 4T1 (breast cancer) **E)** and MC-38 (colon cancer) co-cultures are shown. **F)** The surface marker expression of human CD14^+^ cell co-cultures with A549 (lung cancer), **G)** MDA-MB-231 (short MDA, breast cancer), **H)** and PANC-1 (pancreatic cancer). **F–H)** Cell percentages of CD11c^+^ cell populations are shown. **C–H)** 10 kPa and 20 kPa were stiffened by addition of CaCl_2_ and soft controls were not stiffened alginate/collagen hydrogels. All experiments were performed using three biological repeats per technical repeat and at least two technical repeats in total. One representative technical repeat is shown in each graph. Three biological repeats per condition were used for the statistical analysis. **A+B)** All statistical differences are depicted as the Benjamini-Hochberg procedure corrected p values: *p < 0.05; **p < 0.0001. **C–H)** All p values are shown: *p < 0.05; **p < 0.01; ***p < 0.005; ****p < 0.0001. All error bars are shown as SEM.

Next, it was investigated if cancer cells would induce different phenotypes in co-cultures. For this, B16F10 melanoma cells were added at the start of the culture and CD11c was used to distinguish DCs from melanoma cells. Surprisingly, CD11c^+^/CD86^+^, CD11c^+^/CD14^+^ and CD11c^+^/TLR7^+^ cell populations were almost completely abolished in 2D co-cultures (B16F10), while there was no change to CD11c^+^/CD317^+^ cell populations (Figure 3b).

Next, development of co-cultures in 3D was investigated. In 3D melanoma co-cultures CD11c^+^/CD14^+^ cells increased in 20 kPa gels while CD11c^+^/CD317^+^ cell populations remained unchanged along with CD11c^+^/STAT5p^+^ populations. By contrast, CD11c^+^/CD86^+^ cells decreased on stiffer substrates (Figure 3c). To determine if this was a general mechanism, it was necessary to confirm these results with two other cancer cell types. For this purpose, 3D co-cultures with breast cancer cells (4T1, Figure 3d) and colon cancer cells (MC38, Figure 3e) were established. In both cultures CD11c^+^/CD317^+^ and CD11c^+^/STAT5p^+^ decreased with stiffness, while CD11c^+^/CD86^+^ populations were only present on soft controls. Finally, CD11c^+^/CD14^+^ populations increased with stiffness. This suggests that some cancer cells can prevent CD11c^+^/CD317^+^ from developing at high stiffnesses, which would be beneficial in evading the anti-tumor pDC phenotype altogether. Ultimately, high stiffness suppressed cDC development and favored progenitors developing into CD14^+^ populations.

Finally, 3D co-culture experiments using human cells were performed to validate the murine data. Instead of using bone marrow–derived cells, CD14^+^ cells were isolated from the blood of six healthy donors. Cultures with lung cancer (A549), pancreatic cancer (PANC-1), and breast cancer (MDA-MB-231) were set up and revealed similar results to the murine cultures regarding CD11c^+^/CD86^+^ cell populations (Figures 3f–h). Notably, CD11c^+^/CD317^+^ were not lost on higher stiffnesses but did not exceed 15%. These results mirror the murine results, whereby most cells did not have a cDC or pDC phenotype in higher stiffnesses in cancer co-cultures. Surprisingly, CD11c^+^/CD14^+^ cell populations drastically decreased in higher stiffnesses. It was hypothesized that the initial myeloid progenitors were further developed compared to the murine BM cultures and, thus, might have differentiated into another phenotype, such as a mature macrophage phenotype. To test this hypothesis CD11c^+^/CD16^+^ cell populations were investigated, as CD16^+^ cell populations increased on stiff substrates in 2D. However, CD11c^+^/CD16^+^ cell populations did not increase at all or not to a cell population larger than 5% in higher stiffness gels.

### Tumor-associated macrophage marker expression increases with stiffness

Since most myeloid cells in human 3D cancer co-cultures were neither cDC, pDC nor basic macrophages it was hypothesized that most cells would express TAM markers. One of the best established TAM markers is CD206 (5). Indeed, CD206^+^ cell populations increased with stiffness independent of stimulation in 2D murine monocultures (Figure 4a). Additionally, *VEGF* expression and VEGFA production was increased on higher stiffnesses in 2D (Figure 4b), along with *PTGS2* expression (Figure 4c). Furthermore, out of the seven MMPs that reached mRNA level around or above 1000 reads, five MMPs were significantly increased on stiffer substrates (Figure 4d+e). Out of the seven highest expressed MMPs, *MMP9* was the only gene which was more highly expressed on soft substrates (Figure 4f), while *MMP3* expression levels were not significantly different (Supplementary Figure 1e). In human cancer co-cultures, CD11c^+^/CD206^+^ populations increased with stiffness for lung and pancreatic cancer co-cultures and reached at least 20% across all cancer types in higher stiffnesses (Figure 4g). Furthermore, human MMP13 production increased in 10 kPa compared to soft controls in co-cultures across all cancer types (Figure 4h).

**Figure 4:**
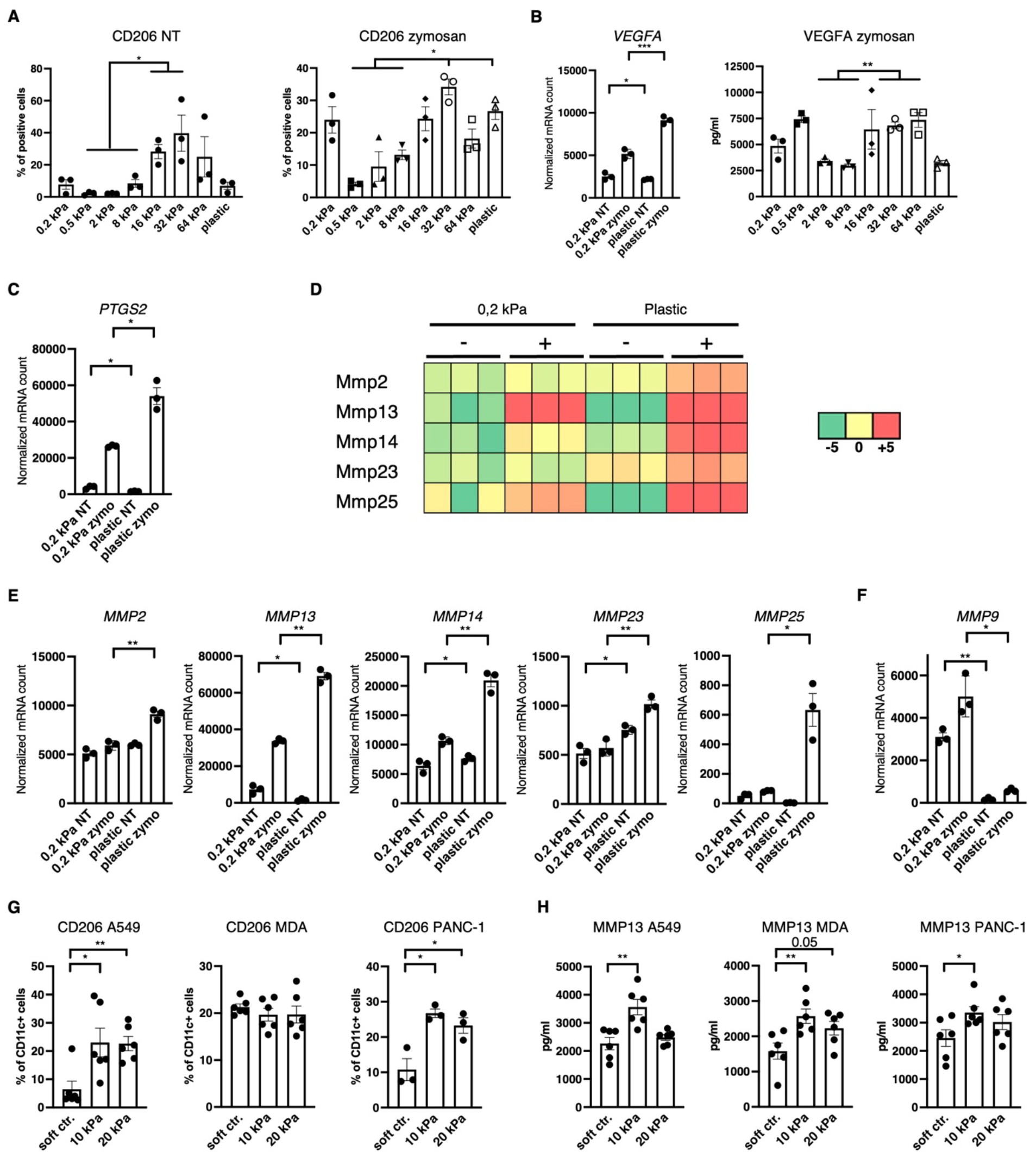
Increased stiffness promotes tumor-associated macrophage marker expression. **A–F)** Data are derived from murine 2D monocultures. **A)** CD206^+^ cell populations of resting cells (NT, not treated) and zymosan-stimulated cells (24 h) were measured via flow cytometry. **B)** Normalized *VEGFA* mRNA counts measured during RNAseq and VEGFA production measured via ELISA from zymosan-stimulated (24 h) cells are shown (zymo=zymosan) **C)** Normalized *PTGS2* (protein also known as COX2) mRNA counts measured during RNAseq are shown. **D)** Heatmap depicting expression level of five of the seven highest expressed MMPs of DCs cultured on 0.2 kPa hydrogels (+/-zymosan) and DCs cultured on plastic (+/-zymosan). **E+F)** Normalized *MMP* mRNA counts measured during RNAseq. **G)** CD206 expression of CD11c^+^ cells measured via flow cytometry and **H)** MMP13 production measured via ELISA of human CD14^+^ cell co-cultures. **G+H)** 10 kPa and 20 kPa were stiffened by addition of CaCl_2_ and soft controls were not stiffened alginate/collagen hydrogels. All experiments, except RNAseq, were performed with three biological repeats per technical repeat and at least two technical repeats in total. RNAseq was performed with one technical repeat. One representative technical repeat is shown in each graph. Three biological repeats per condition were used for the statistical analysis. All statistical differences are shown: *p < 0.05; **p < 0.01; ***p < 0.005; ****p < 0.0001. All error bars are shown as SEM.

### Akt regulates DC and macrophage phenotypes in a species-specific way

TAM polarization towards M2 has been suggested to be regulated via the PI3K/Akt pathway (22-24) and Akt phosphorylation is reportedly negatively regulated by β2-integrins in DCs (12). Akt was thus hypothesized to be a potential regulator of the DC phenotype to either increase expression of pDC and cDC surface marker in higher stiffness or reduce TAM marker in stiffer substrates. To target the Akt pathway Akt Inhibitor IV (Calbiochem) was used, which targets a kinase upstream of Akt and downstream of PI3-K. In murine 3D cancer co-cultures, Akt inhibition indeed led to a decrease in CD11c^+^/CD206^+^ cell populations across all cancer types in stiffer gels (Figure 5a). Akt inhibition did not impact the CD11c^+^/CD86^+^ population percentages in murine cultures (Figure 5b). In human 3D cancer co-cultures, however, Akt inhibition did not impact CD11c^+^/CD206^+^ population (Figure 5C), while a notable reduction to MMP13 production was observed only in pancreatic cancer co-cultures (Supplementary Figure 1f). Instead, Akt inhibition led to a significant increase in CD11c^+^/CD86^+^ populations across all stiffnesses and cancer types. This indicates that Akt is an important regulator of the myeloid phenotype, but its specific regulation mechanism is species-dependent.

**Figure 5:**
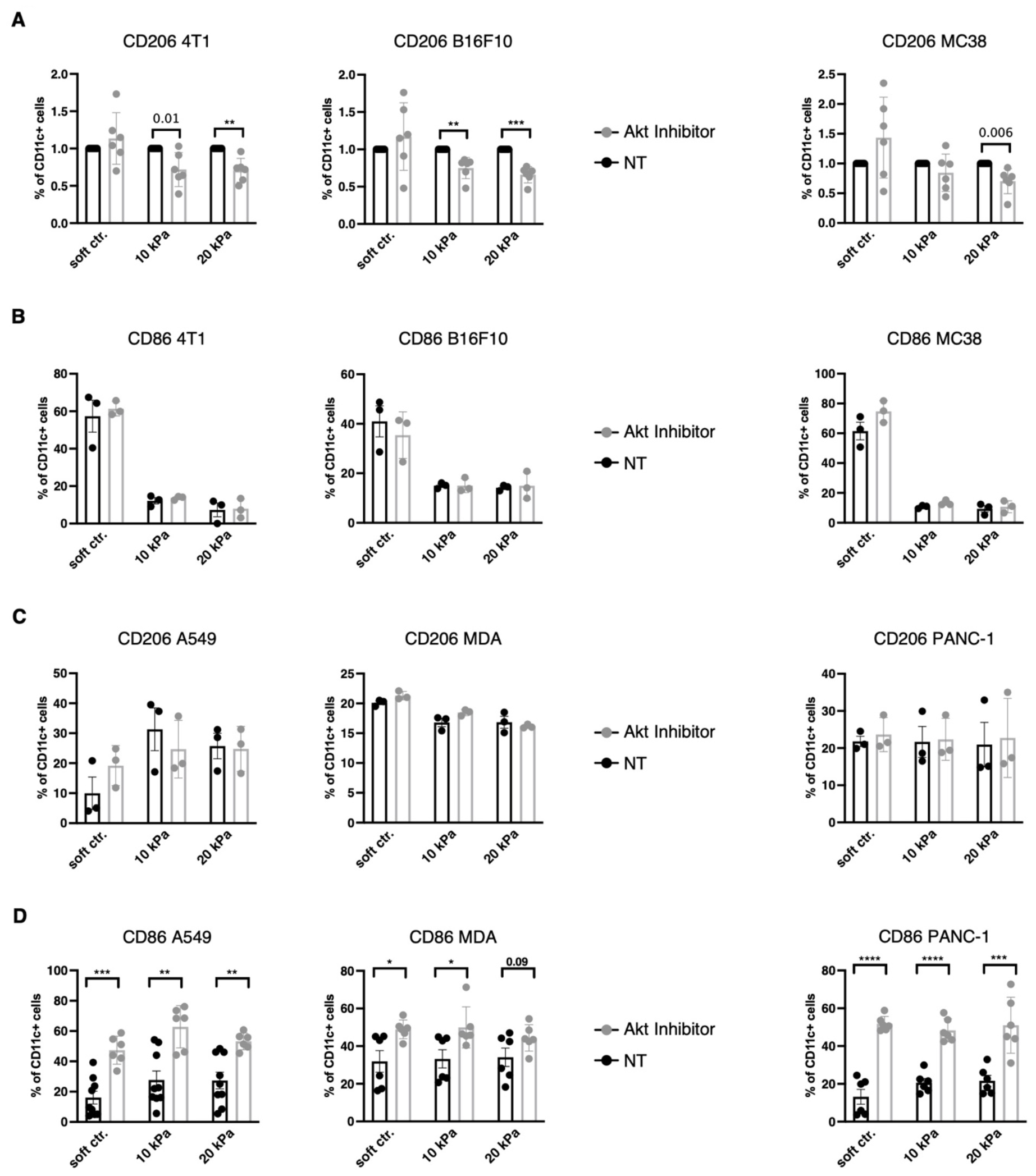
Akt regulates different DC/macrophage markers in human compared to in mice. **A)** For all graphs, the individual control (NT, not treated) percentages were set to 1 for each stiffness, upon which the corresponding fold change was calculated. **A)** CD206 and **B)** CD86 expressions of CD11c^+^ cells were measured via flow cytometry. **C)** CD206 and **D)** CD86 expressions of human CD11c^+^ cells were measured via flow cytometry. **A-D)** Akt was inhibited with 1.2 μM Akt Inhibitor IV (Cayman) for 24 h. All experiments were performed with three biological repeats per technical repeat and at least two technical repeats in total. For **A)** and **D)** all repeats are shown and were used for the statistical analysis (n = 6 per condition). For **B)** and **C)** one representative technical repeat is shown and three biological repeats per condition were used for the statistical analysis. **A–D)** 10 kPa and 20 kPa were stiffened by addition of CaCl_2_ and soft controls were not stiffened alginate/collagen hydrogels. All p values are shown: *p < 0.05; **p < 0.01; ***p < 0.005; ****p < 0.0001. All error bars are shown as SEM.

## Discussion

The aim of this study was to map the impact of tissue stiffness on DC subset development. Previous reports indicated that integrins, especially β2-integrins, restrict the mature cDC phenotype (12), while a mature cDC phenotype can be induced by blocking overall adhesion of differentiated WT BMDC or plating them on soft hydrogels overnight (12, 13). However, such phenotypical changes are dynamic and a cDC phenotype that was induced by blocking adhesion begins to reverse six hours after adhesion is re-established (13). What has not been investigated thus far is the outcome of the long-term differentiation on different substrate stiffnesses. The results in this paper show that such cultures result in distinct cDC phenotypes on soft matrices and pDC phenotypes on stiff substrates. While the pDC markers CD317, Siglec-H, and TLR7 were all upregulated, B220 was barely expressed. Furthermore, interferon RNA was not detected in cells cultured on stiff substrates. This indicates that high stiffness merely contributes to the pDC phenotype, while other chemical or mechanical signals are required to achieve a functional pDC phenotype.

Nonetheless, this has implications for chronic diseases in which stiffness increases. In this report, this mechanism was investigated in the context of cancer. This study’s results indicate that, in some cancers like melanoma and lung cancer, these pDC-like cells might arise due to stiffness cues, while in other cancers like colon cancer other mechanisms might suppress the expression of pDC markers altogether. What is common across the cancer types investigated, was an increase in macrophage markers, especially TAM markers in humans. While TAMs are well established as promoting tumor progression (5), a CD14+ DC subtype has also been previously linked to a immunosuppressive tumor microenvironment (21). Therefore, stiffness could be a hidden contributor to DC dysregulation in cancer.

In this study 3D co-cultures were used, specifically collagen/alginate hydrogels, which were originally established by Elysse C. Filipe et al. (25). To current knowledge, this culture system is the only 3D culture system that allows A) long-term (7+ days) culture without toxicity, B) cell adhesion, C) cell retrieval after culture and D) the generation of matrices of up to 20 kPa without changing the chemical ratio. In terms of immunological studies, one of the biggest drawbacks of this system is the component alginate. Alginate has been previously shown to induce LPS-like immune responses in macrophages (26). Because alginate is a polysaccharide, it is highly likely recognized by CLRs and, for this reason, zymosan was not used for further stimulation on 3D cultures as these cultures were already believed to be stimulated. Additionally, the use of alginate renders the use of these hydrogels *in vivo* inherently flawed.

Nevertheless, this culture system was utilized to identify a potential mechanism via which myeloid cells, specifically DCs, become dysregulated and twisted to support tumor progression by secreting VEGFA and MMPs. Initial attempts to target this mechanism intracellularly were partially successful but revealed an even higher level of complexity. This study confirms other reports of the PI3K/Akt signaling pathway regulating tumor-associated macrophage markers such as CD206 in murine macrophages (23). However, this regulation does not seem to work in human cells in the same way. In human DCs, Akt signaling has been shown to be downstream of CD80/CD86 signaling (27). To current knowledge, the results presented in this study are the first to demonstrate that Akt inhibition led to an increase in CD86 expression in human CD11c+ cells. Combined these two studies’ results could indicate that CD86 expression is regulated in a feedback loop in human DCs.

Ultimately, this study found that stiffness governs DC and macrophage phenotypes, a mechanism which possibly contributes to DC and macrophage dysregulation in cancer. How exactly these phenotypes are regulated intracellularly remains unclear, but the mechanism seems to be species-specific. Similar species-specific differences might exist in general cell signaling and contribute to most drugs identified on the bench failing during clinical trials (6). Nevertheless and in any context, Akt was identified as an important regulator of DC and macrophage surface marker expression. This is relevant, because both reduction of TAM markers and increase of CD86 expression might be beneficial for increased patient survival and targeting Akt could be a viable strategy to achieve this.

## Materials and Methods

### Mice

C57BL/6 mice were purchased from CLEA Japan, Inc (Tokyo, Japan). All mice were maintained in a filtered-air laminar-flow enclosure and given standard laboratory food and water *ad libitum*. All animal protocols were approved by the Animal Care and Use Committee of the Research Institute for Microbial Diseases, Osaka University, Japan (Biken-AP-R03-17-1).

### Cell culture

Murine DCs were differentiated from isolated bone marrow via culture in RPMI + 10% FCS 1% Pen/Strep and 1% Amino Acids containing 20 ng/ml GMCSF (Biolegend, #576306) on silicone hydrogels with different stiffnesses: 0.2, 0.5, 2, 8, 16, 32, and 64 kPa (Advanced BioMatrix, # 5190-7EA). Hydrogels were coated with 100 ug/ml collagen I (Nippi, #ASC-1-100-100) or 1:100 diluted Matrigel (Falcon, #354234) according to Advanced BioMatrix’s instructions.

For 3D cell cultures cells were embedded in alginate gels modified from the protocol by Elysse C. Filipe et al. (25). Briefly, four volumes of a 2% alginate (NovaMatrix, #P-1408-24 or Funakoshi, #KIM-19130-100) solution in 0.9% saline solution were mixed with two volumes of 3 mg/ml collagen I stock solution (Nippi, #ASC-1-100-100). One volume of 0.9% saline, NaOH (3.9 mM final concentration, Sigma Aldrich, #S8045), and CaCO_3_ (5 mM final concentration, Sigma Aldrich, #C4830) was added, along with bone marrow cell suspension and cancer cell suspension, where applicable. Polymerization was initiated with D-(+)-Gluconic acid δ-lactone (4.2 mg/ml final concentration, Sigma Aldrich #G4750). 1ml of gel per well was cast in 24 well plates. Gels were left to polymerize overnight at 37°C and then stiffened with a calcium chloride (Sigma Aldrich, #C1016) solution of 25 mM for 10 kPa or 50 mM for 20 kPa for 30 min at 37°C. For soft stiffness conditions, no calcium chloride was added to the polymerized gels. At the end of the culture, gels were digested with 100 μg/ml alginate lyase (Merck, #A1603-100mg) and 1 mg/ml collagenase (Sigma, #C5138) for 2 h at 37°C.

For co-cultures with cancer cell lines, the following murine cancer cell lines were used: B16F10 (melanoma, from Riken Bioresource Center), MC-38 (colon cancer), and 4T1 (breast cancer). Murine colon and breast cancer cell lines were a kind gift from the Sakaguchi lab. For human cancer cell co-cultures, the following human cell lines were used: A549 (lung cancer, ATCC, #CCL-185), MDA-MB-231 (breast cancer, ATCC, #HTB-26), and PANC-1 (pancreatic cancer, ATCC, #CRL-1469). Cancer cells were added at the beginning of the culture at a 1:10 ratio with bone marrow cells or human CD14^+^ monocytes.

### Human monocytes

Human CD14+ monocytes were isolated from the peripheral blood of healthy donors using lymphocyte separation solution based gradient centrifugation (Nacalai Tesque, #20828-44) followed by magnetic bead sorting utilizing anti-human CD14+ antibodies (Miltenyi Biotech, #130-050-201). CD14+ monocytes were cultured in RPMI-1640 media supplemented with 10% FCS, 0.1 mM non-essential amino acids (Gibco, #11140-050), 1 mM sodium pyruvate (Nacalai Tesque, #06977-34), 1% Pen/Strep, 0.1% Amphotericin B (Thermo Fisher Scientific, #15290026), 10 ng/ml hGM-CSF (Peprotech, #576306), and 10 ng/ml hIL-4 (Peprotech, #574004) for eight days. Donors had given informed consent in accordance with the Declaration of Helsinki prior to blood donation. Donors consisted of four females and two males aged 25– 39. The institutional review board of Osaka University approved the blood draw protocols for healthy individuals (approval number 29-4-10).

### Stimulation and inhibitor

Dendritic cells were stimulated or inhibited on day 9 of the culture for 24 h. Cell stimulation was done with 100 μg/ml zymosan (Sigma, #Z4250-1G), 100 μg/ml depleted zymosan (Nacalai, #59008-84), 20 ng/ml LPS (Sigma, L4516), 10 ug/ml trehalose 6,6’-dimycolate (Sigma, T3034) or 10 ug/ml β-glucosylceramide (Avanti Polar Lipids, #860549P-5mg). To inhibit Akt, cell cultures were treated with 1.2 μM Akt Inhibitor IV (Cayman, #15569) for 24 h.

### RNA sequencing

For RNA sequencing, cells were directly lysed on the hydrogels with ice-cold Trizol (Thermo, #15596026). Stimulated cells were treated 20 h before lysis. Lysates were collected and submitted to the NGS service of the University of Osaka. Briefly, total RNA was extracted using a RNeasy Mini kit (QIAGEN) and library preparation was performed using a TruSeq-stranded mRNA sample prep kit (Illumina, San Diego, CA, USA) according to the manufacturer’s instructions. Whole-transcriptome sequencing was then performed with the RNA samples using the Illumina HiSeq 2500, HiSeq 3000 or NovaSeq 6000 platforms in the 75-or 101-base single-end mode. Sequenced reads were mapped to the mouse reference genome sequences (mm10) using the TopHat software, version 2.1.1 (John Hopkin’s University, Baltimore, MA, USA). The number of fragments per kilobase of exon per million mapped fragments (FPKMs) was calculated using the Cufflinks (GitHub) software, version 2.2.1. Unsupervised k-means clustering analysis was performed via the integrated Differential Expression and Pathway analysis website (http://bioinformatics.sdstate.edu/idep/). RNA-seq data was deposited at the Sequence Read Archive under the Accession number: PRJNA1082503.

### Flow cytometry

After detaching the cells, FC stain was performed (1:50, BD Pharmingen #553142). The following antibodies were used at a concentration of 1:100: CD14 (BioLegend, #123312, clone Sa14-2), CD86 (BioLegend, #105012, clone GL-1), Langerin (BioLegend, #BL144203, clone 4C7), CD317 (BioLegend, #127108, clone 129C1), CD16 (BioLegend, #158008, clone S17014E), CD205 (BioLegend, #138214, clone NLDC-145), CD11b (BioLegend, #101207, clone M1/70), CD11c (BP Pharmingen, #5538C1, clone HL3 and BioLegend, #117310, clone N418), TLR7 (BioLegend, #160004, clone A94B10), Siglec-H (BioLegend, #129603, clone 551), CD319 (BioLegend #152003, clone 4G2), STAT5p (BD Pharmingen, #612567, clone 47/Stat5), LY6C (BD Pharmingen, #553104, clone AL-21), STAT4p (Invitrogen, #17-9044-41, clone 4LURPIE), CD80 (BD Pharmingen, #553768, clone 16-10A1), Ikaros (BioLegend, #653303, clone 2A9/Ikaros), STAT1p (BioLegend, #686403, clone A15158B), STAT3p (BioLegend, #651020, clone 13A3-1), B220 (eBioscience, #11-0452-85, clone RA3-6B2) CD206 (BioLegend, #141727, clone C068C2), I-A/I-E (BioLegend, #107621, clone M5/114.15.2), hCD16 (BD Pharmingen, #560918, clone 3G8), hCD14 (BD Pharmingen, #561029, clone M5E2), hCD317 (BD Biosciences, #566381, clone Y129), hCD11c (BD Pharmingen, #560999, clone B-ly6), hCD86 (BioLegend, #305412, clone IT2.2), and hCD206 (BioLegend, #321136, clone 15-2). Dead cells were stained with 2 μg/ml propidium iodide (Sigma, #P4170) or 7AAD (1:100, BioLegend, #420404).

Intracellular staining was done after permeabilization/fixation with the eBioscience Foxp3 transcription factor staining buffer set (Invitrogen, #00-5523-00) according to the manufacturer’s instructions. Samples were measured with an Attune NxT (Thermo Fisher Scientific, #A24858) and analyzed via FlowJo (TreeStar).

### ELISA

The following ELISA kits were used according to the manufacturer’s instructions: TNF (BD Biosciences, #558534), IL-6 (BD Biosciences, #555240), IL-2 (BD Biosciences, #555148), VEGF_165_ (Peprotech, #900-K99), and human MMP13 (R&D, #DY511).

### Statistics

All experiments were performed with three biological repeats and were repeated at least twice (technical repeats, one technical repeat consisted of three biological repeats), except for the RNA sequencing and unless there was no statistically significant difference observed (such as to compare stimulations) to reduce the number of animals used. To calculate the statistical significance and generate the graphs, Graph Pad Prism 9 was used. Unpaired, two-tailed Student’s t-tests or Mann–Whitney tests were used unless otherwise indicated. All p values are shown as *p < 0.05, **p < 0.01, ***p < 0.005, and ****p < 0.0001. For treatments and comparisons shown as fold change Benjamini-Hochberg procedure corrected p values were used (q), for these the values are depicted as follows: *p < 0.05 and **p < 0.0001.

## Acknowledgments

CG would like to thank the IFReC Kishimoto Foundation (for a Kishimoto Fellowship) and the Japanese Society for the Promotion of Science for awarding a Grant in Aid for Young Researchers (23K14541). CG also would like to thank Charles Schutt for advice and guidance, Vanessa Fuller for language editing, and Yamin Qian for the support and advice regarding the cancer co-culture experiments.

## Authorship Contributions

CG designed, performed, and analyzed all experiments unless otherwise specified (RNAseq). CG wrote the paper and created all of the figures.

