## Supplementary Figure 1 for "Stiffness regulates dendritic cell and macrophage subtype development and increased stiffness induces a tumor-associated macrophage phenotype in cancer co-cultures"

### Supplementary Figure 1

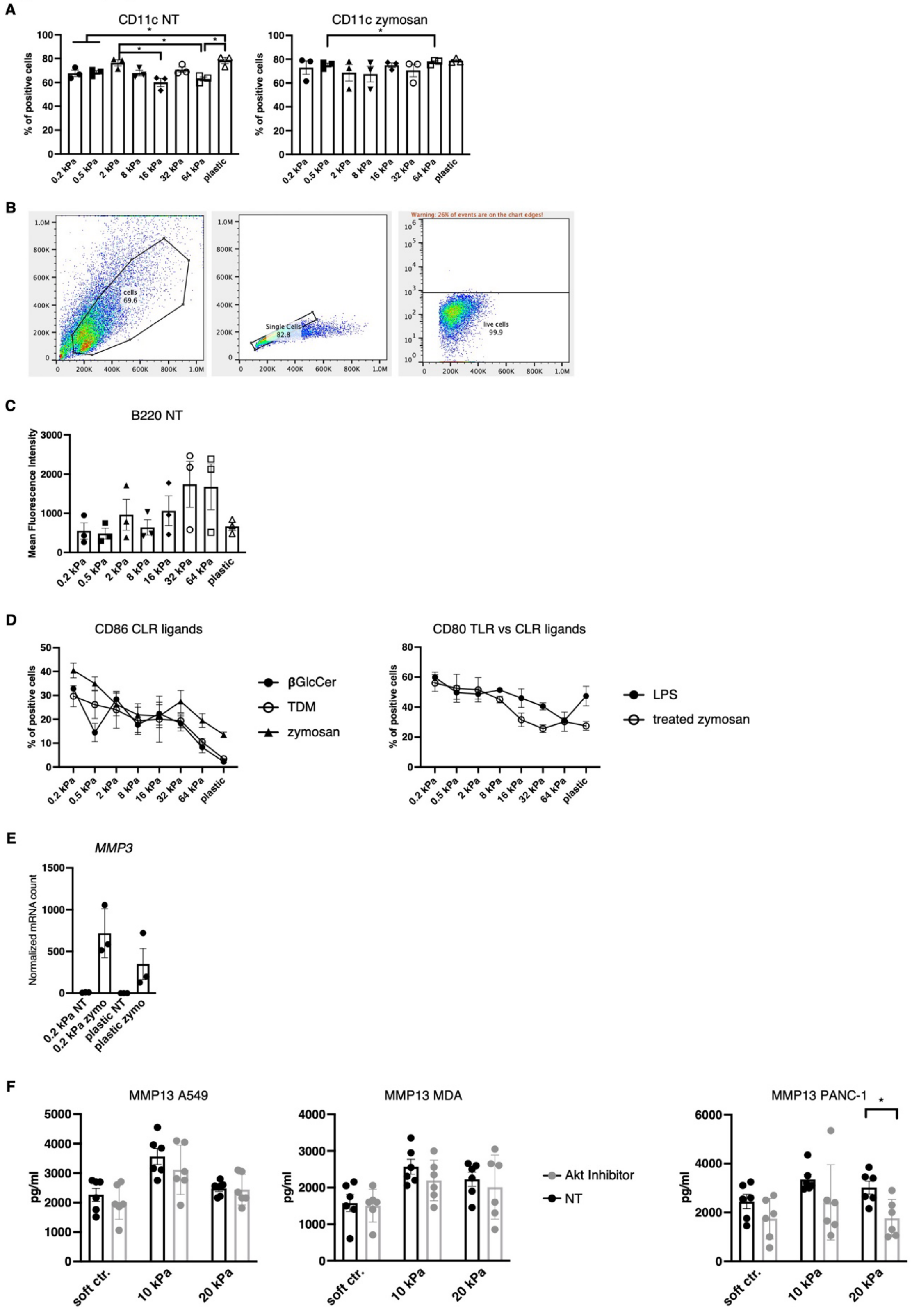

**Supplementary Figure 1)**

**A)** CD11c$^+$ cell populations were measured via flow cytometry on resting cells (NT, not treated) and zymosan-stimulated cells. Percentages of all living cells are shown. **B)** Gating strategy for flow cytometry measurements. **C)** The mean fluorescence intensity (MFI) of B220 from resting BMDC cultures (NT, not treated) measured via flow cytometry. Percentages of all living cells are shown. **D)** CD86 expressions as an example of surface marker expression after stimulation with Mincle DAMP ligand (β-glucosylceramide [beta GlcCer]) and PAMP (trehalose 6,6′ -dimycolate [TDM]) along with Dectin-1 and TLR4 ligand zymosan (left). On the right, stimulation with LPS (TLRs) compared to treated zymosan (Dectin-1 only) is depicted, for which CD80 expression was measured. **E)** Normalized MMP3 mRNA counts measured during RNAseq are shown (NT, not treated; zymo = zymosan). RNAseq was performed with murine BMDCs cultured on 2D hydrogels and compared to plastic. **F)** MMP13 production was measured via ELISA of human CD14$^+$ cell co-cultures. The statistical differences are shown as $^*p < 0.05$; 10 kPa and 20 kPa were stiffened by the addition of $CaCl_2$ and soft controls were not stiffened alginate/collagen hydrogels. Akt was inhibited with 1.2 μM Akt Inhibitor IV (Cayman) for 24 h. **A–F)** Experiments were performed with three biological repeats per technical repeat and at least two technical repeats in total (except RNAseq).
